# Efficient and reliable measles reprogramming platform for the generation of human iPSC

**DOI:** 10.1101/2025.10.10.677451

**Authors:** Saksham Vashistha, Sarrianna Hoffer, Jessica Dade, Spencer Majerus, Austin Royster, Carter Caya, Brenna Sharp, Patricia Devaux

## Abstract

We previously established the Measles virus (MeV) vector for reprogramming somatic cells into induced pluripotent stem cells (iPSCs); however, efficiency was limited to 0.2%. Here, we present the next generation that reprograms with an average efficiency of up to 2.3% and with similar or superior efficiency to the Sendai system. Twenty of the ninety iPSC isolated clones were amplified and analyzed. All clones showed a strong induction of endogenous pluripotency-associated markers SSEA-4, TRA-1-81, TRA-1-60, and NANOG. Analysis of the N mRNA transcript over passages showed rapid elimination of the vector from the iPSC at or before passage four. The pluripotency propensity was further analyzed using spontaneous and guided differentiation into three germ layers. All clones showed similar ability to differentiate into hematopoietic, pancreatic, and neuronal progenitor cells. In conclusion, this study shows that this new MeV reprogramming vector has a comparable or higher reprogramming efficiency than currently available systems and offers faster vector elimination from iPSCs. It uses a lower multiplicity of transduction as it uses a single vector versus multiple vectors. Finally, MeV produces iPSCs that can differentiate into multiple cell types. This makes MeV an efficient reprogramming platform for iPSC generation from patient samples.

## Introduction

Nearly twenty years ago, induced pluripotent stem cells (iPSCs) were first introduced as a promising technology in the field of stem cell research ^1-3^,. By subjecting somatic cells to a reprogramming process, researchers in Japan were able to return these cells back toward a state of pluripotency. This process involved the delivery of four reprogramming factors (RFs) – OCT4, SOX2, KLF4, and cMYC or L-MYC (OSKM) into somatic cells to induce a pluripotent state ^2^. The resulting iPSC showed similar morphology and properties to those of Embryonic Stem Cells (ESCs) ^1^, ^4^ and presented a novel source of stem cells, which contrasts to ESCs, which require serious ethical considerations before being used ^5^. Furthermore, there is a growing need for the rapid development and production of autologous iPSCs from patients for individualized therapy, since the use of allogeneic iPSCs or ESCs requires long-term use of immunosuppressive therapy to avoid rejection of cellular therapies. While the development of off-the-shelf universal iPSCs is slowly developing, the safety of these cells in patients remains to be determined ^6^.

Various integrating and non-integrating methods have been developed to deliver the necessary factors to a cell that induce reprogramming to an iPSC ^7^, ^8^. DNA viral vector methods include adenoviral and adeno-associated viral vectors ^9-12^,; however, the efficiency of these vectors is either low (≤ 0.001%) or largely unsuccessful in adult human fibroblasts and peripheral blood mononuclear cells (PBMCs) ^9-13^,. Several non-viral reprogramming methods have been successful as well. This includes transduction of proteins ^14^, DNA minicircles ^15^, transposons ^16^-^18^, replicating episomal vectors based on Epstein-Barr virus (EBV) ^19-21^, or messenger RNA transfection ^22^. While each of these methods has been successful in reprogramming somatic cells, most of them have been shown to be unreliable at reprogramming blood cells ^23^. The Sendai vector (SeV) has held a leading role in the reprogramming field ^24^, but its reliance on three viral vectors and high MOI transduction often leads to a lingering expression of the vector beyond 12 passages in the iPSC, slowing down the stabilization of the iPSC clones, and producing partially reprogrammed iPSCs ^25^. Due to these limitations, there is a need for an efficient, reliable, and safe system of iPSC production that can be translated quickly into the clinical setting.

We have previously developed a single-cycle vector based on the vaccine strain of the measles virus (MeV) and showed that we could express the four reprogramming factors in a single genome of MeV and reprogram adult human fibroblasts (AHFs) at an efficiency of 0.04% ^26^. In later studies, we improved the efficiency of the system to 0.2% ^27^; however, this efficiency was still much lower than the Sendai or mRNA systems, which report efficiencies between 0.5-1.4%, or 0.6-4.4%, respectively, depending on the cell type ^28^. Here, we introduce the next generation of measles-based reprogramming vector and demonstrate its ability to reprogram 6 different AHFs with an efficiency between 0.2 and 2.2% depending on the fibroblast. We isolated and characterized over 20 MeV-derived iPSCs and showed the presence of all pluripotency markers in all clones, rapid vector elimination by passage 4, and high genomic stability. All MeV-derived iPSCs demonstrated a strong propensity for spontaneous and guided differentiation into the three germ layers, as well as the ability to easily differentiate into pancreatic, neuronal, and hematopoietic progenitor cells. Lastly, we conducted a comparative analysis between SeV and MeV on the same AHFs used in the study and demonstrated that MeV reprogramming vector significantly outperforms SeV in 50% of fibroblasts tested and matched SeV for the remaining AHFs. This new single-component MeV reprogramming system not only achieves high efficiency, but also prevents the formation of partially reprogrammed iPSCs, improving clone differentiation and stability for effective translation to clinical applications.

## Results

### Production of an improved and efficient reprogramming MeV vector

We have previously produced a single-cycle MeV expressing the four RFs, MV(O)(SK)(M), to produce iPSCs. MV(O)(SK)(M) expresses the *OCT4* and *SOX2-KLF4* bicistronic (SK) genes in two independent additional transcription units (ATU) instead of the MeV hemagglutinin (*H*) gene and the *c-MYC* gene in an additional transcription unit (ATU) after the *H* gene (Figure 1A, top genome ^27^). This vector was fully based on the background of the Moraten vaccine strain of MeV. We modified this vector by switching the MeV phosphoprotein (*P)* gene to a *P* gene from a wild-type strain of measles virus, Ichinose-B (IC-B)^29^, MVP^WT^(O)(SK)(M) (Figure 1A, bottom genome). To track the vectors, the *GFP* gene was inserted into the ATU after the *P* gene. The new vector MVP^WT^(O)(SK)(M) showed comparable growth kinetics to the previously published vector MV(O)(SK)(M) and replicated to a maximum titer of about 10^6^ by 48h, confirming that vector propagation remains unaltered by the substitution of the P gene (Fig. 1B).

**Figure 1:**
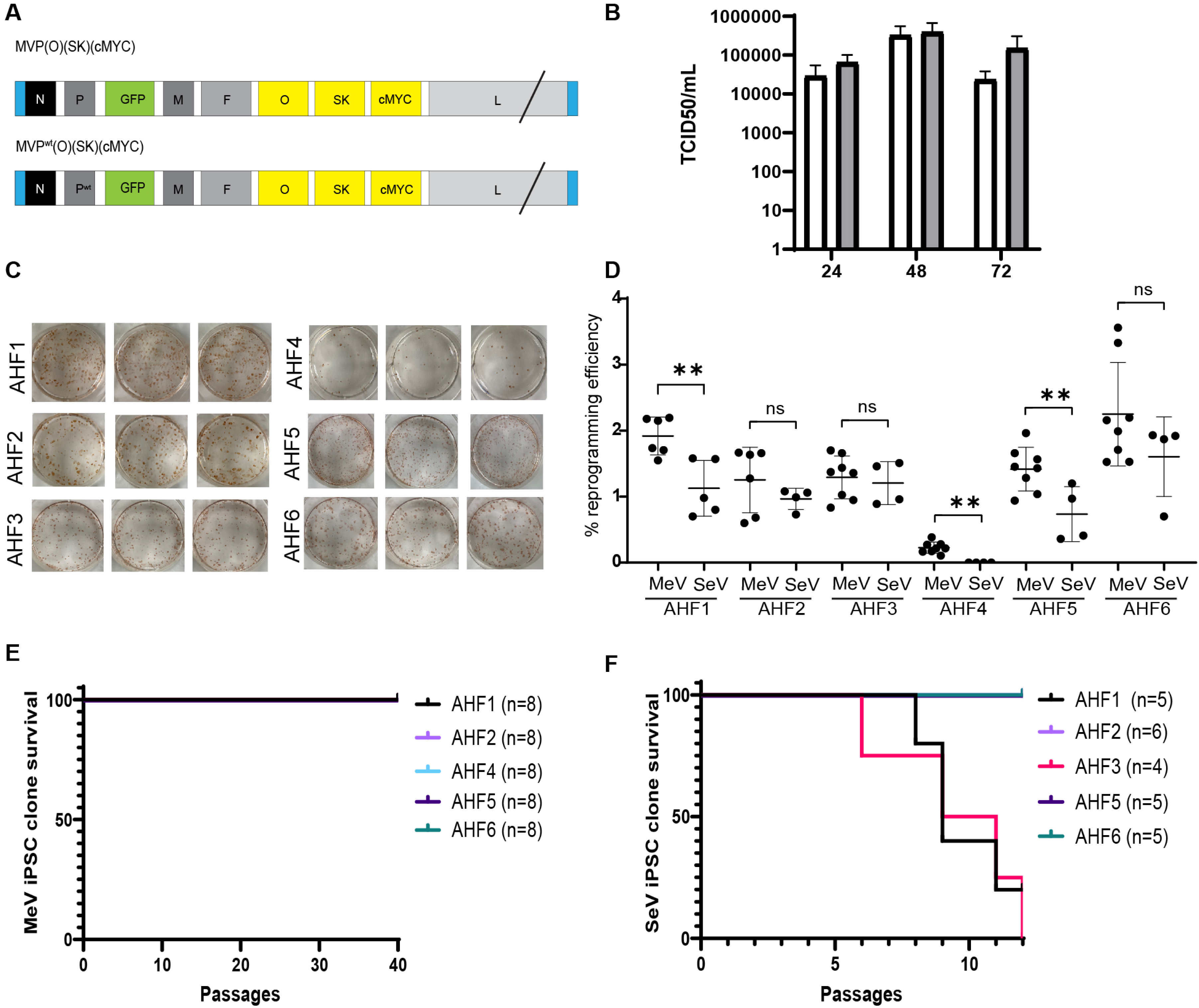
Reprogramming with MVP^WT^(O)(SK)(M) leads to efficient generation of iPSC clones and is more efficient than SeV. (A). Structure of the MeV reprogramming vector MVP^WT^(O)(SK)(M) and parental vector MV(O)(SK)(M). (B) Titers of cell-associated and released virus produced upon infection of Vero-H2 cells with MVP^WT^(O)(SK)(M) (white) and MVP^Vac^(O)(SK)(M) (^27^, grey) vectors at 24, 48, and 72h post-infection. Error bars represent SD from three individual experiments. An unpaired t-test was used (all non-significant, not labeled). (C) Representative NovaRED staining of one reprogramming for AHF1 to AHF6. One reprogramming from a 12-well is split into 3 x 6-well. Each red dot represents one iPSC clone stained by TRA-1.60 antibody. (D) Average reprogramming efficiency by MVP^WT^(O)(SK)(M) or SeV system (Cytotune 2.0) as indicated for AHF1 to AHF6, n=4-8. Data presented as mean ± SD. An unpaired t-test was used to compare the vectors for each AHF (** = p < 0.005, ns= not significant). (E) Survival curve of MeV-derived iPSC clone over 40 passages for AFH1, AHF2, AHF4, AHF5 and AHF6. n=8. (F) Survival curve of SeV-derived iPSC clone over 12 passages for AFH1, AHF2, AHF3, AHF5 and AHF6. n=4-6.

### MVP^WT^(O)(SK)(M) vector can efficiently reprogram adult human fibroblasts into iPSC-like cells

We had previously reported the efficiency of reprogramming for the MV(O)(SK)(M) to be 0.04% in AHF ^27^. To determine the reprogramming efficiency of the MVP^WT^(O)(SK)(M) vector, six AHF (3 males and 3 females) were reprogrammed with the new vector (Table 1). Six to eight reprogramming experiments were performed for each fibroblast following the established protocol previously described ^27^. Briefly, 5×10^4^ fibroblasts were transduced with MVP^WT^(O)(SK)(M) vector at an MOI of 0.5, spinoculated for 1h at 250g and incubated overnight at 37 °C. Media was changed every day thereafter, and cells were split into three Matrigel-coated wells of a 6-well plate on day 6, then transitioned to reprogramming media.

Nine to twelve days post-transduction, GFP-positive, sharp-edge, flat, and tightly packed iPSC-like colonies started to appear in the replated Matrigel-coated 6-well plate, and multiple iPSC-like clones were found in the transduced cells by day 15. On day 22, staining with TRA-1-60, a marker of pluripotency and detection with NovaRED was performed to determine the number of iPSC clones per well. Figure 1C shows representative staining from one reprogramming experiment of each AHF with MVP^WT^(O)(SK)(M). The number of iPSC clones was tallied by two independent analysts to ensure the accuracy of the results. The reprogramming efficiency reached an average of 0.23% to 2.25% depending on the AHF (Fig. 1D). All AHFs, except one, generated an average of 646 to 1370 iPSC clones, and only AHF4 produced a lower number of iPSC clones (average of 113). We compared the reprogramming efficiency of MVP^WT^(O)(SK)(M) to the commercially available SeV system (Cytotune 2.0). The same AHFs were reprogrammed, and the average efficiency for the SeV system was either equivalent or statistically lower than the efficiency observed with MeV (Fig.1D). Interestingly, while MeV was able to reprogram AHF4, SeV was unable to produce any iPSC clones for this specific AHF. Single iPSC clones derived from MeV reprogramming were picked and expanded. Eighty MeV iPSCs (four clones from four independent reprogramming) were derived from five of the six AHF. The iPSC clones picked from AHF3, were removed from the study due to a technical mistake during propagation. All other MeV-iPSC clones survived, and despite limited differentiation in the early passages, stabilized, and remained stable for subsequent passages up to passage 40 (Fig. 1E). Twenty iPSC clones, 4 clones per AHF (one clone from the four independent reprogramming) were further analyzed for vector elimination, pluripotency markers, differentiation propensity, fingerprinting, karyotypes, and tested for mycoplasma. Since the SeV system is a well-established technology, only four to six iPSC clones from the five AHF that reprogrammed were picked, for a total of 25 iPSCs, and amplified to passage 12. As AHF4 did not reprogram entirely, no clone could be propagated. Out of the 25 clones, only 17 could be propagated to P12. All clones from AHF3 and four out of the five iPSC clones from AHF1 did not stabilize and ended up differentiating (between P5 and P8), indicating that most of the clones generated from these two AHFs were only partially reprogrammed (Fig. 1F). Finally, the SeV clones were not further investigated for elimination of the vector from the iPSC, pluripotency and multilineage differentiation propensity, as these characterizations are well established for Sendai-derived clones ^23^ and the goal of this study is to validate the new MVP^WT^(O)(SK)(M) vector reprogramming technology and the iPSC clones derived from this vector.

### After passage 4, the MeV reprogramming vector is cleared in over 95% of the clones and 100% after passage 5

Twenty clones were analyzed for MeV vector elimination over the first seven passages following reprogramming. To determine the MeV vector genome’s persistence in derived iPSC clones, we targeted the most highly expressed MeV gene – nucleoprotein (*N*) – for detection using quantitative PCR. Using a previously established MeV standard curve for the qPCR, we determined that the qPCR detection limit was 4 molecules of *N* ^26^. We showed that 10% (2/20), 40% (8/20), 65% (13/20), 95% (19/20), 100% (20/20), and 100% (20/20) of the clones eliminated the MeV reprogramming vector by passage 1 (P1), 2 (P2), 3 (P3), 4 (P4), 5 (P5) and 6/7 (P6/7), respectively. One positive result and two positive results for passages P4 and P5, respectively, were counted as qPCR contamination, because they were negative in previous passages. Therefore, *N* was only detected in one iPSC clone at P4 (Fig. 2A and B), indicating fast clearance of the MeV vector after reprogramming. We attribute this result to the system’s initial low MOI of transduction and the fast multiplication of the iPSCs, diluting the vector over each passage.

**Figure 2.**
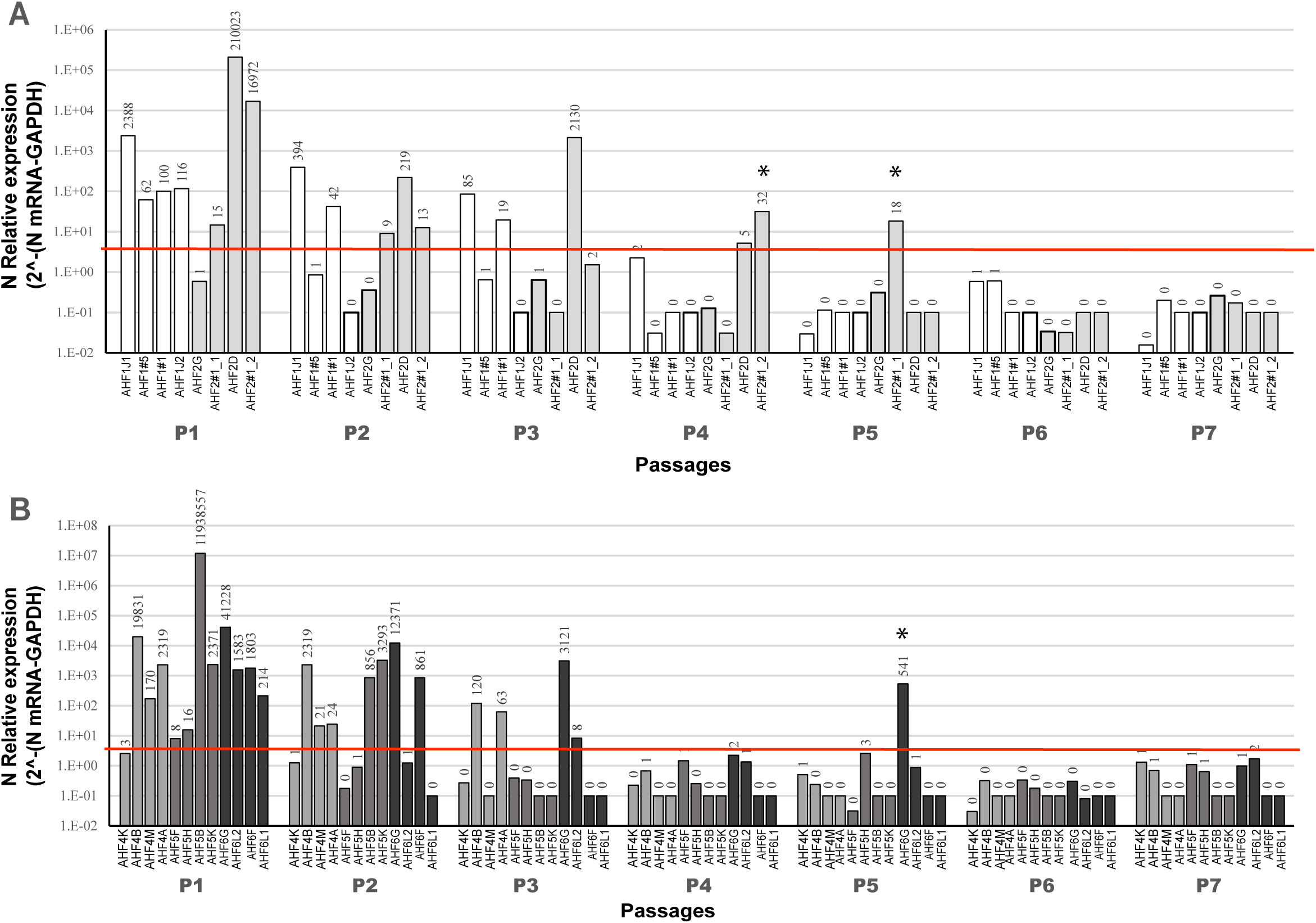
Rapid clearance of MeV vector during iPSC expansion. (A) qPCR analysis of N-gene transcripts across serial passages of four iPSC clones derived from male AHF1 and AHF2. (B) qPCR analysis of *N*-gene transcripts across serial passages of four iPSC clones derived from female AHF4, AHF5 and AHF6. Number = number of *N* copies. The red line marks the limit of detection. * Represent clones with PCR contamination.

### All MeV-derived iPSC clones displayed characteristics of human iPSCs

The 20 iPSC clones were characterized for the expression of pluripotency markers between passages 10-12. All 20 iPSC clones expressed pluripotency-associated markers, SSEA-4, TRA-1-60, TRA-1-81, OCT4, SOX2, and NANOG, without distinction of being raised from male or female AHF. Figure 3A shows a representative set of 4 iPSC clones, two derived from male and 2 from female fibroblasts and the other iPSC clones are shown in supplemental figure S1A. To confirm that the MeV-derived iPSC-like clones are pluripotent, we further characterized them for their propensity to differentiate into the three germ layers: endoderm, ectoderm, and mesoderm. All 20 MeV-derived iPSCs could form Embryoid Bodies (EBs) when cultured in suspension and differentiate spontaneously into ectoderm, mesoderm, or endoderm, as shown by the expression of beta III tubulin, CD31 or FOXA2, respectively (Fig. 3B and Supplemental Fig. S1B).

**Figure 3.**
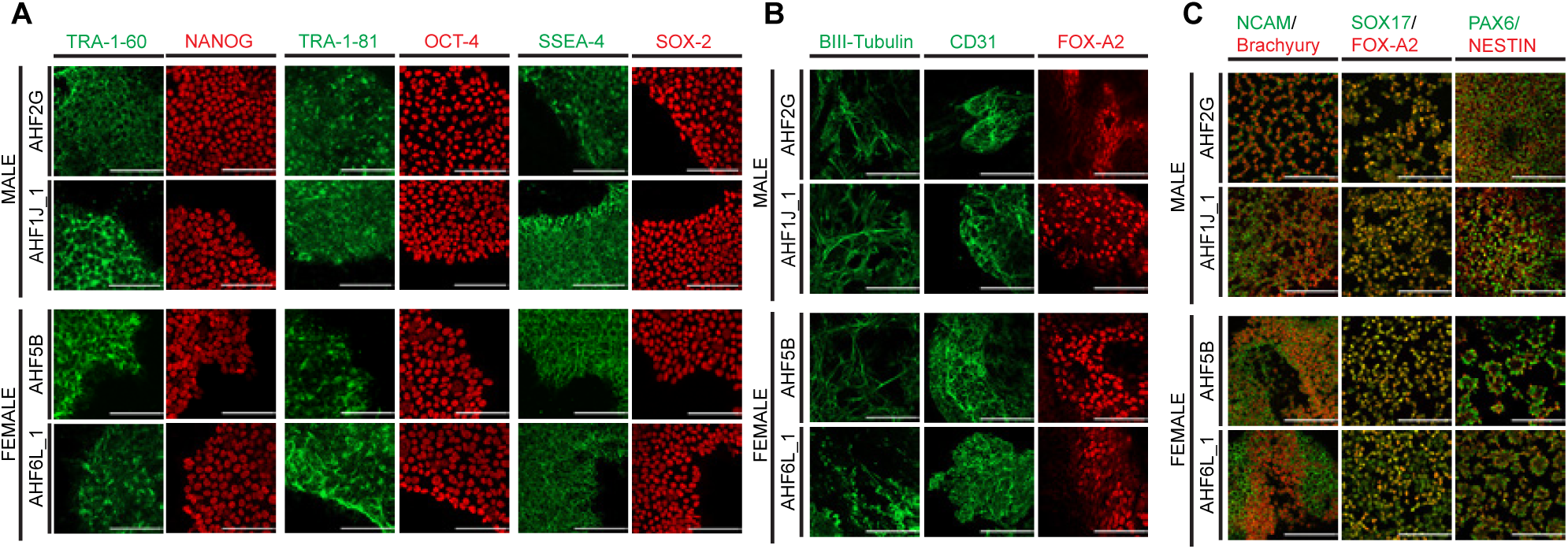
MeV-derived iPSCs express pluripotency markers and can differentiate into the three germ layers. (A) Confocal microscopy imaging of four representative MeV-derived iPSCs with dual immunostaining for pluripotency markers TRA-1-60, TRA1-81, SSEA-4, NANOG, OCT4, and SOX2. Scale bar: 100 μm. (B) Confocal microscopy imaging of spontaneous differentiation of four representative MeV-derived iPSCs into embryoid bodies (EBs). Cells were immunostained for EB markers βIII-tubulin, CD31, and FOX-A2. Scale bar: 100 μm. (C) Guided trilineage differentiation into mesoderm, endoderm, and ectoderm germ tissues of four representative MeV-derived iPSCs. Cells were immunostained for associated lineage markers using NCAM (green)/Brachyury (red) for Mesoderm, SOX 17 (green)/FOXA2 (red) for Endoderm and PAX6 (green)/NESTIN (red) for Ectoderm lineages. Scale bar: 100 µm

However, while all the iPSC clones appeared to spontaneously differentiate equally into ectoderm and mesoderm, it is important to note that for a few clones, the endodermal lineage was not strongly represented, as shown by visibly low SOX17 and FOXA2 expression (AHF2#1_2, AHF5K, and AHF6L_2). Following guided differentiation, all iPSCs tested differentiated into the ectodermal, endodermal, or mesodermal pathways, confirming the multi-lineage differentiation propensity of all MeV-derived iPSCs (Fig. 3C and Supplemental Fig. S1C). From the guided differentiation, we could observe a few clones that had a strong propensity to differentiate into mesodermal tissue (for example, AHF2#1_1, AHF6L1 or AHF5B) or ectodermal tissue (AHF4B, or AHF6L2). The three clones (AHF2#1_2, AHF5K, and AHF6L2) that did not differentiate well into endoderm during EB and spontaneous differentiation did not appear to have difficulties differentiating into endoderm during the directed differentiation protocol (Fig. 3B and C and Supplemental Fig. S1B and C).

### All MeV-derived iPSC clones can differentiate into progenitor cells

We next explored the ability of the 20 iPSCs to differentiate into three different types of progenitor cells: neuronal (NPC), pancreatic and hematopoietic (HSC). The 20 iPSC clones were first subjected to differentiation into NPC using the STEMdiff Neural Induction Medium and SMADi for 7 days and seeded into Matrigel-coated 4-well chamber slides for 2 to 4 days before immunostaining (Fig. 4A and Supplemental Fig. S2). To confirm NPC differentiation, the cells were stained with antibodies against PAX6, SOX1, and NESTIN. Figure 4A shows representative images of the differentiated iPSC clones into NPC after 12-14 days of differentiation and stained with the three NPC markers. All iPSC clones differentiated into NPC and expressed a significant staining with the three markers, SOX1, PAX6, and NESTIN, indicating that all the clones analyzed can progress through the ectodermal pathway and further into NPCs.

**Figure 4.**
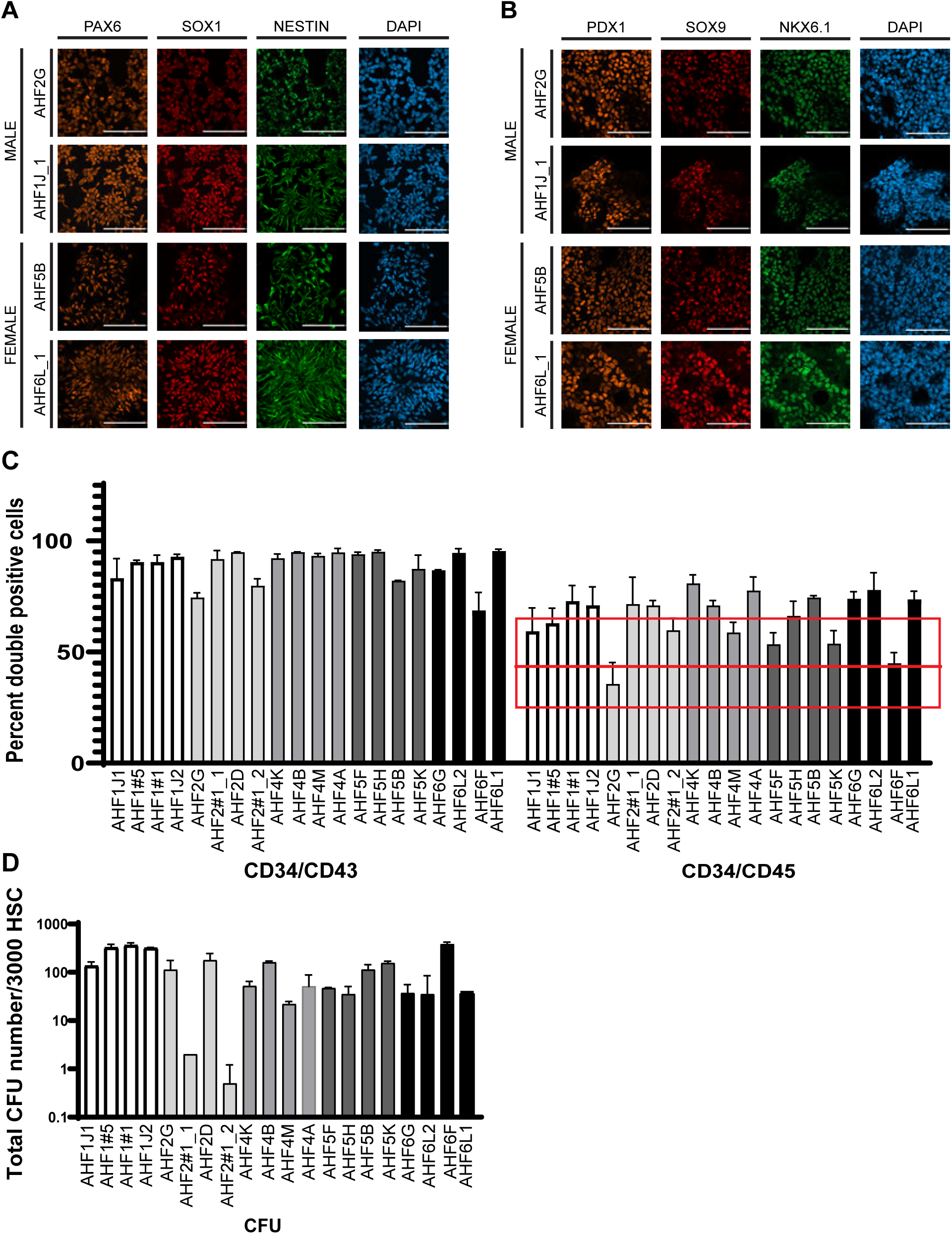
MeV-derived iPSC can differentiate into multiple progenitor cell types. (A) Confocal microscopy imaging of neuronal progenitor differentiation from four MeV-derived iPSCs. Cells were stained with PAX6 (orange), SOX1 (red), NESTIN (green), and DAPI (blue). Scale bar: 100 µm (B) Confocal microscopy imaging of pancreatic progenitor differentiation from four MeV-derived iPSCs. Cells were immunostained for PDX1 (orange), SOX9 (red), NKX6.1 (green), and DAPI (blue). Scale bar: 100 µm. (C) Percent of CD34/CD43 (left) and CD34/CD45 (right) double-positive cells after hematopoietic stem cell differentiation of the 20 MeV-derived iPSC clones. The red box corresponds to the predicted differentiation window expected (25%-65%), and the red line corresponds to the average (43%) indicated by the manufacturer. Average of three experiments. (D) Percent of colony-forming units (CFU) tallied after the CFU assay of the 20 MeV-derived iPSC clones. Average CFU from three experiments.

Next, 20 iPSC clones were seeded in a Matrigel-coated 4-well chamber slide and subjected to differentiation into pancreatic progenitors using the STEMdiff pancreatic progenitor differentiation kit for 14 days before immunostaining (Fig. 4B and Supplemental Fig. S2B). The pancreatic differentiation was confirmed by staining with antibodies against PDX1, SOX9, and NKX6.1. Figure 4B shows representative images of the differentiated iPSC clones into pancreatic progenitor cells and shows that all the iPSC clones expressed PDX1, NKX6.1, and SOX9, following pancreatic differentiation, indicating that all the clones analyzed can not only differentiate into NPCs, but also through the endodermal pathway and into pancreatic progenitors.

Next, the 20 iPSC clones were seeded in Matrigel-coated 12-well plates and subjected to differentiation into HSC progenitors using the STEMdiff Hematopoietic Kit for 12 days. On day 12, the cells were stained for markers of HSC differentiation, CD34, CD43, and CD45 and then analyzed by flow cytometry. Figure 4C summarizes the results of HSC differentiation and shows the percentage of double-positive CD34 /CD45 and CD34^+^/CD43^+^ cells for the 20 iPSC clones. The CD34 /CD45 cells represent a population of hematopoietic stem and progenitor cells that are often isolated as a functional progenitor population capable of colony formation (CFU assays). The double-positive CD34^+^/CD43^+^ are a population of early hematopoietic progenitors. In iPSC/ESC differentiation, CD34^+^/CD43^+^ cells are considered definitive hematopoietic progenitors with the potential to give rise to multiple blood lineages (myeloid, lymphoid, erythroid). This figure shows that all iPSC clones differentiated into HSC, with the percentage of CD34^+^/CD43^+^ and CD34 /CD45 cells ranging between 55-96% and 35-75%, respectively. According to the manufacturer’s kit, differentiation typically yields between 25-65% CD34 /CD45 progenitor cells (average 43%, red box and red line, respectively). All 20 iPSC clones matched or exceeded the predicted differentiation efficiency provided by the supplier (Fig. 4C). Only one clone, AHF2G, did not reach the average of 43%, with only around 35% of CD34 /CD45 cells. The CD34 /CD45 cells include progenitor cells that have the capacity to form hematopoietic colonies in the colony-forming unit (CFU) assay. Thus, to confirm the ability of the HSC to further differentiate into the hematopoietic lineage, a CFU assay was performed. At day 10 of the differentiation, 3000 HSC were isolated, seeded and differentiated further for 14 days utilizing MethoCult™ SF H4636. All 20 iPSC clones up to this point could generate terminal hematopoietic cells as demonstrated by countable colony-forming units (Fig. 4D) and were further characterized for specific colony types, including BFU, CFU-GM, CFU-GEMM, and others (not shown), indicating that all the clones analyzed can progress through the mesodermal pathway into HSC progenitors and terminal hematopoietic cells.

### Validation of the genomic stability and identity of the MeV-derived iPSC clones

Finally, to validate the genomic stability and identity of the iPSC clones, karyotyping and fingerprinting analysis were performed. Karyotyping of the starting adult human fibroblast at P5 was performed, and all AHFs had a normal karyotype (Fig. 5, top row). All iPSC clones were tested at passage 25 or higher, and 19/20 showed normal diploid karyotypes (Fig. 5, iPSC section). Only one iPSC (AHF6F) had an 18q deletion in 2 of the 20 cells that were analyzed (Fig. 5, red box), indicating that the vast majority of the MeV-derived iPSCs did not go through major chromosomal rearrangement over the first 25 passages. Analysis by fingerprinting confirmed the identity of all the iPSC clones analyzed in this study with their respective AHF (not shown). Mycoplasma testing was performed to validate that all data obtained in this study were from mycoplasma-free iPSC clones, and all clones tested negative (not shown).

**Figure 5.**
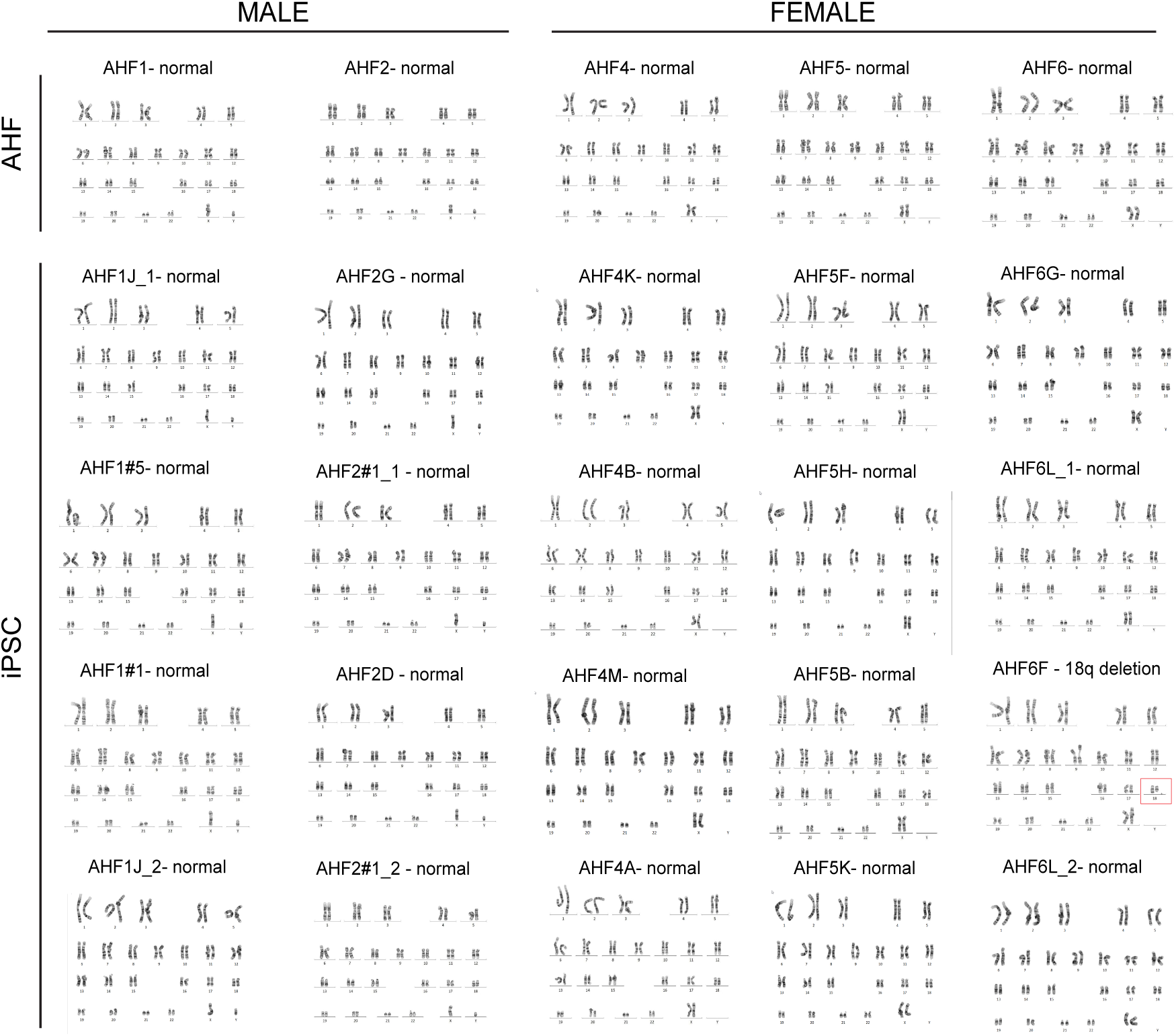
MeV-derived iPSCs show chromosomal stability over passages. Representative G-banding chromosome analysis of parental AHFs (top row) and the 20 MeV-derived iPSC clones.

## Discussion

In 2019, we developed a proof-of-concept measles reprogramming system based on the Food and Drug Administration’s (FDA) MeV vaccine that was able to produce iPSCs from neonatal fibroblasts, but this vector had a poor reprogramming efficiency (0.002%) and did not reprogram AHF. Later, we reported an improved vector with an efficiency of 0.04% in AHF; however, this vector still underperformed compared to the most used reprogramming systems such as SeV, reprogramming AHF at around 1%. In this study, we report on the latest generation of MeV vectors that reprogram with an efficiency between 0.2% to 2.5% in AHF. When we performed a direct efficiency comparison between MeV and the established SeV system, we found that MeV was able to outperform SeV for the reprogramming of three AHFs - AHF1, AHF4, and AHF5 - and matched SeV in the three other AHFs. Most notably, the SeV system was unable to reprogram AHF4 entirely, while the MeV system was able to achieve a measurable efficiency of around 0.2% and produce stable iPSC clones that were successfully analyzed. We believe that this difference in reprogramming performance, particularly in refractory cell lines like AHF4, is likely due to the MeV’s vector design – a single vector containing all four reprogramming factors as well as its natural tropism for human cells. In contrast, as SeV is a murine virus, there is a possibility that SeV entry and/or transcription could be affected in some human cells. Thus, MeV’s strong performance across multiple different fibroblasts, as well as producing stable iPSC with refractory lines, highlights the system’s reliability in reprogramming.

In addition to its efficiency, the MeV reprogramming platform confers several key advantages over other non-integrating tools, such as Sendai ^30-32^,, the Episomal system ^19^, and even mRNA ^33^. MeV uses a single component, which not only bolsters the ease and efficiency of transduction but also eliminates the possibility of producing partially reprogrammed iPSCs, as all the RFs are present in every transduced cell. With multi-component technologies, it is possible that transduced or transfected cells express sub-optimal concentrations of one or more RFs, despite a high MOI or high DNA transfection, leading to the production of partially reprogrammed cells or unstable iPSC clones. In this study, we have found that some of the randomly picked SeV clones could not be propagated for two of the AHF lines (AHF1 and AHF3) indicating that there may have been generation of more unstable iPSC clones in some AHFs than in others. Furthermore, the protocol of the Cytotune 2.0 tool recommends selecting SeV-derived iPSC clones after performing a live staining with TRA-1-60 or TRA-1-81. In contrast, for MeV-derived iPSC clones, all clones were picked solely based on their iPSC-like shape, and all clones propagated as iPSCs up to passage 40, indicating that most, if not all, MeV-derived iPSCs are fully reprogrammed and that our MeV system and protocol eliminate these additional live staining steps.

Another advantage is the MeV reprogramming platform’s rapid elimination from the iPSC. We showed that the MeV is eliminated in all clones tested by passage 4. While we use an anti-measles antiviral between day 18 and 21, which does help with the elimination of the vector, the residual vector is primarily eliminated by dilution through iPSC division and passages, as we have shown previously ^26^. This rapid elimination contrasts with other systems, like Sendai or Episomal DNA, which takes up to 9 to 12 passages or over 20 passages to eliminate the reprogramming components from their respective iPSC clones ^23^. In 2022, Charlesworth et al. described a new Sendai system, still based on three components, in which they modified the KLF4 vector and introduced a miR367 target sequence. This modification led to the rapid elimination of SeV genomes in iPSCs, decreasing the limit of detection to 34 days from the onset of transduction instead of 42 days. This was after culturing the cells for 14 days at 32 °C, followed by 20 days at 38 °C, ^31^, ^32^. Using the MeV vector, the clones are picked between days 23 and 25 and then passaged every 4 days, meaning that the first iPSC clones (10%) were vector-free by day 26 to 28 (2/20, P1) if not before. The percentage of vector-free iPSC increase after each passage to 40% on day 30 to 32 (8/20, P2), 65% on day 34 to 36 (13/20, P3), 95% on day 38 to 40 (19/20, P4) and 100% on day 42 to 44 (20/20, P5) and this is achieved without additional modification of the vector or shifting the cells to different temperature conditions during reprogramming. Moreover, this faster elimination is likely due to the lower initial MOI used for the transduction of the AHF. As a single vector, MeV only requires a low MOI of 0.5 compared to SeV, which requires three vectors and uses a higher MOI for each vector, 5:5:3, for a total MOI of 13. Rapid vector clearance is not only critical for faster use of iPSC in the laboratory but also highlights improvements in the technology’s safety profile by eliminating residual vector components quickly, opening the door for potential GMP applications.

We had established a proof-of-concept vector using the attenuated vaccine strain of MeV for the reprogramming of iPSC ^26^, ^27^. However, these systems conferred a low efficiency of reprogramming in AHF. In this study, we replaced the vaccine *P*-protein with the wild-type *P* protein (*P^wt^*), which significantly improved the reprogramming efficiency of the vector from 0.04% to up to 2.2% in AHF and to 0.2% in AHF difficult to reprogram. We hypothesize that this improvement could be linked to one of the functions of the proteins encoded by the *P* gene. The MeV *P* gene encodes for two additional proteins, the *C* and *V* proteins, which are known to interfere with the innate immune response ^34-40^,, and there is evidence that the wild-type proteins are better at controlling the innate immune response ^41^, ^42^. This could affect the reprogramming process. It was shown that the activation of innate immune signaling through toll-like receptors (TLR) or RIG-1-like receptors (RLR) is a possible contributor to the efficient reprogramming in adult fibroblasts ^22^, ^43^. The TLR and RLR pathways can mediate changes in epigenetic plasticity to facilitate reprogramming ^44^, ^45^. In human cells, the RLR family is represented by RIG-1, MDA5, and LGP2, which each exert their effects by signaling through the adaptor protein interferon-beta promoter stimulator 1 (IPS-1) ^46^. The *V* and *C* proteins of MeV have been reported to directly interfere with and affect the activation of these proteins ^47^. Therefore, there is a possibility that the interaction of the wild-type proteins, and not the vaccine’s attenuated counterparts, with one of these three proteins is responsible for the increase in reprogramming efficiency.

Finally, MeV-derived iPSC showed complete expression of pluripotency markers as well as differentiation into the three germ layers. Furthermore, MeV-derived iPSC demonstrated robust tissue-specific differentiation, expressing markers of pancreatic, neuronal, and hematopoietic progenitors. Hematopoietic differentiation was functionally validated with CFU assays, which revealed not only a high colony count but also a broad spectrum of CFU subtypes. The presence of CFU colonies suggests that MeV-derived HSC are highly multipotent and capable of giving rise to a variety of hematopoietic lineages. This extensive analysis indicated that the MeV-derived iPSCs exhibit high genomic stability and robust multipotency, as evidenced by strong and complete differentiation into various target tissues.

In conclusion, while iPSCs hold transformative potential in regenerative medicine, current non-integrating reprogramming tools still suffer from slow clearance, partial reprogramming, and technical complexity. This study demonstrates that the MeV reprogramming platform addresses several of these critical barriers by offering a highly reliable, efficient, and user-friendly reprogramming system to produce stable and rapid vector-free human iPSCs.

## Materials and Methods

### Cells

Six de-identified human adult skin fibroblasts from healthy donors (three males and three females) were obtained from the Mayo Clinic Center for Regenerative Biotherapeutic Biotrust under IRB # 22-001576. These samples have already been collected and archived by the Mayo Clinic Center for Regenerative Medicine Biotrust (under IRB #13-007298, Dr. Wigle). The samples were made available to the investigators as de-identified samples. The only information provided to the research team was the age and gender of the donor. Fibroblasts were maintained in DMEM with 10% ES-FBS (Life Technologies, Carlsbad, CA, USA), 0.1mM non-essential amino acids (Corning Mediatech, Manassas, VA, USA), and 1% Penicillin and Streptomycin (P/S, Corning Mediatech,).

Rescue-H2 cells (helper 293-3-46-H2) and Vero-H2 cells were described previously ^48^ and were maintained in DMEM with 10% FBS (Life Technologies), and 1% P/S (DMEM-10 medium). G418 at 1.2mg/mL was added to the media of the rescue cell line. All cell lines above were cultured in a humidified atmosphere with 5% CO2 at 37°C.

The iPSC clones were maintained in mTeSR1 medium (STEMCELL Technologies, Vancouver, Canada), containing 1% P/S on 12 well-matrigel-coated plates (Corning Mediatech). When 80– 90% confluency was reached, iPSCs were split using EZ-Lift (Sigma Aldrich, St. Louis, MO, USA) following the manufacturer’s instructions and propagated as clumps in new Matrigel-coated plates. iPSCs were frozen in CryoStor CS10 (Sigma Aldrich). The iPSC clones were cultured in a humidified atmosphere with 5% CO2 at 37°C.

### Vector rescue and production

The rescue of recombinant MVP^wt^(O)(SK)(M) was performed according to previously published procedures ^48^ using the Rescue-H2 cells. In brief, Rescue-H2 cells were transfected using calcium phosphate precipitation (ProFection kit, Promega, Madison, WI, USA) with the MV genome plasmid and MV polymerase plasmid (pEMCLa). The rescue-H2 cells were then overlaid on Vero-H2 cells, three days post transfection. Green fluorescent protein (GFP) expression was used to monitor the appearance of infectious centers, and single viruses were isolated and propagated on new Vero-H2 cells. For MVP^wt^(O)(SK)(M) stock production, Vero-H2 cells were infected at a multiplicity of infection (MOI) of 0.05 in OptiMEM (Life Technologies) for 2 h at 37°C. DMEM-10 medium was added, and the cells were incubated overnight at 37 °C. In the morning, the cells were transferred to 32°C until 95% of the cells expressed GFP. Cell culture media was removed, and cells were scraped in 1ml OptiMEM. Vector particles were released by two freeze-thaw cycles. Titers of virus stocks were determined by 50% end-point dilution (tissue culture infectious dose 50, or TCID50) on Vero-H2 cells using the Spearman–Kärber method ^49^.

### Growth curves

VeroH2 cells (4×10^5^cells) were infected with MeV vector at an MOI of 0.05 in OptiMEM. After 2 h, the vector was removed, washed, and 1ml of DMEM 10% was added. Cells were collected at 24-, 48-, and 72-hour post-infection, along with the media. The samples were later titered using TCID50 titration Assay ^49^.

### Measles vector Reprogramming

Adult human fibroblast cells (5 × 10^4^) were seeded on matrigel-coated plates and transduced with MVP^WT^(O)(SK)(M)^19^ at 0.5 MOI in OptiMEM. Cell and virus were subjected to spinoculation at 250g for 1 h at 25°C, then the inoculum was left overnight at 37°C on top of the cells. The day after, the cells were washed once with PBS, and DMEM-10%ES-FBS-NEAA-PSP (media1) was added. The media was changed every day until day 6, when the cells were split using TrypLE into 3 wells of a 6-well plate coated with Matrigel. Cells were slowly transferred to reprogramming media containing 80% Nutristem (Sartorious, Ann Arbor, MI, USA), 20 ng/ml of basic human FGF (STEMCELL Technologies), 1% P/S, and 20% mTeSR1 medium (STEMCELL Technologies) from day 6 to 9, (25%-75%, 50%-50%, 75%-25%, and 100%-0% reprogramming/BJ media % ratio, respectively) and reprogramming media was changed every day thereafter. Small molecules (SM); SB431542 (5 μM), PD0325901 (0.2 μM), and Thiazovivin (0.5 μM) (all from Stemgent/Reprocell, Cambridge, MA, USA), were added to the media from day 6 to 14 ^27^, ^50^. From day 18 to 21, 5 μM of AS-136A anti-measles drug (MedChemExpress, Monmouth Junction, NJ, USA; ^51^) was added to the reprogramming media for 3 days, to help eliminate the vector. Around day 20-22, the iPSC clones were picked based on size and morphology and transferred individually to a matrigel-coated 24-well plate for further expansion and analysis. Reprogramming efficiency was calculated as the percent of TRA-1-60-positive iPSC colonies generated divided by the number of input cells. TRA-1-60-positive colonies were visualized with NovaRED HRP substrate (Vector Laboratories, Burlingame, California, USA), according to the manufacturer and the number of clones was tallied by two independent analysts for accuracy.

### Sendai Vector Reprogramming

Sendai reprogramming was performed using CytoTune™-iPS 2.0 Sendai Reprogramming Kit (Invitrogen). 4 x 10^4^ AHF cells were seeded in a 12-well plate two days before transduction of cells. An additional counting well was seeded along with any experimental wells to determine the correct vector transduction. The viral titer was calculated and reported on the manufacturer’s certificate of authentication (COA). At the time of transduction, viral volumes were calculated according to the following calculation: Cells counted x MOI/viral titer. Virus was left in the well overnight at 37°C, then the cells were washed once with PBS, and DMEM-10%-ES-FBS-NEAA-PSP was added the following day. According to the protocol, media was changed on days 2, 4, and 6. At day 7, each well was split using TrypLE into 3 wells of a 6-well plate coated with Matrigel. Following the split, the cells were then media changed daily using Essential 8 media (Thermo Fisher Scientific, Waltham, MA, USA) as recommended by the protocol. At day 25, the iPSC clones were picked based on size and morphology and transferred individually to a matrigel-coated 24-well plate for further expansion. Reprogramming efficiency was calculated as above for the MeV vector.

### RNA collection, Viral gene transcription and N quantitative qPCR

A confluent well (80-90%) of 12-well plate of iPSCs was detached with EZ-Lift, and iPSCs were pelleted by centrifugation at 200g and resuspended in 1ml of Trizol reagent (Thermo Fisher Scientific) and stored at −80°C until RNA extraction was performed. MeV-iPSC cells were collected at passages 1, 2, 3, 4, 5, 6, and 7. Total RNA was isolated with Trizol reagent according to the manufacturer’s protocols. RNA was quantified using a NanoDrop 2000 Spectrophotometer (Thermo Fisher Scientific). RNA to cDNA reverse transcription was performed with the EcoDry TM Premix (Oligo dT) kit (Takara Bio, Shiga, Japan) using 1µg of RNA according to the manufacturer’s instructions. Quantitative PCR was performed with N (Fwd: 5’CGGAGCTAAGAAGGTGGATAAA3’; Rev: 5’CAGTCCAAGAGCAGGATACATAG3’) and GAPDH (IDT PrimeTime Predesigned Primers, Hs.PT.58.40035104, NM_002046) using Bullseye EvaGreen qPCR Master Mix (MidSci, St Louis, MO, USA) with 2µl of cDNA in a final reaction volume of 25 µl. As a positive control, we performed a quantitative PCR on dilution of different amounts of MeV vectors in 1µl of cDNA from 4LV-iPSC clone (40000, 4000, 400, 40, 4, 0.4 molecules of MeV vectors were used). The negative control was 1µl of cDNA from an iPSC derived with four Lentiviral vectors encoding each a reprogramming factor ^26^. Three technical replicates were performed for each sample. Measurements were collected using QuantStudio 7 Pro Real-Time PCR systems (Thermo Fisher Scientific) at 55°C annealing temperature for 40 cycles. Relative gene expression of N and GAPDH mRNAs were compared using 2^(-DΔC_T_) method.

### Immunostaining and confocal microscopy for pluripotency markers

To analyze undifferentiated iPSCs, iPSCs were seeded on Matrigel-coated (Corning) chamber slides (Lab Tek II). Cells were washed with PBS and fixed in PBS-2% PFA for 15 minutes. Cells were then permeabilized in PBS-2% PFA-0.1%Triton X-100 for 15 minutes, washed 3 times with PBS and incubated overnight in a blocking solution (PBS-5% FCS). Overnight incubation with primary antibodies. Primary antibody staining was performed with mouse anti-SSEA-4, TRA1-60, and TRA1-81 (1:100), rabbit anti-OCT4 (1:200), rabbit anti-SOX2 (1:200), and NANOG (1:100) diluted in blocking solution. Corresponding Alexa Fluor 488 anti-mouse or Alexa Fluor 594 anti-rabbit (1:1000) were used for secondary antibody staining for 2h at RT. After five washes with PBS, cells were mounted with Prolong Gold Antifade reagent containing DAPI (4’,6-diamidino-2-phenylindole) (Life Technologies). Confocal microscopy analyses were performed using a Zeiss LSM 980 confocal microscope, followed by analysis with Zen blue software (Zeiss). All antibodies used are described in Table S2.

### Spontaneous differentiation assay

MV-derived iPSC clones were detached using TrypLE and seeded in low-adhesion 96-well plates at 16,000 iPSC/well in iPSC medium, spun at 300g for 10 minutes and cultured for 10 days in suspension to form Embryoid Bodies (EBs). After 10 days, EBs were moved to matrigel-coated chamber slides (LABTECK^R^ II) for attachment and cultured in DMEM 20% ES-FCS for an additional 10 days to allow for differentiation. Differentiated cells were fixed in PBS-4%PFA for 15 minutes at RT, permeabilized for 30 minutes, blocked and immunostained with either rabbit anti-FOXA2 (1:100) for endoderm, chicken anti-ß-III tubulin (1:1000) for ectoderm, and mouse anti-CD31 (1:50) for mesoderm diluted in blocking solution overnight at 4°C. After three PBS washes, secondary antibodies Alexa Fluor 594 anti-rabbit (1:1000), FITC anti-chicken (1:500), Alexa Flour 488 anti-mouse (1:1000) were used on corresponding primary antibodies as described above. Confocal microscopy analysis was performed as described above. All antibodies used are described in Table S2.

### Guided differentiation assay

Guided differentiation assay was performed using STEMdiff^TM^ Trilineage Differentiation Kit (Stemcell Technologies) according to the manufacturer’s directions. iPSCs were seeded on a Matrigel-coated 24-well glass-bottom plate according to the density defined in the manufacturer’s protocol. Differentiation was performed for 5 days for mesoderm and endoderm and 7 days for ectoderm. Differentiated cells were fixed for 15 minutes at RT in PFA 2%, permeabilized for 15 minutes in PBS-PFA2%-Triton 0.1%, blocked in PBS-FBS5%, and immunostained using markers suggested by the manufacturer’s protocol. Rabbit anti-Nestin (1:500) and mouse anti-Pax-6 (1:100) were used for Ectoderm staining. Rabbit anti-FoxA2 (1:100) and mouse anti-SOX17 1:100) were used for Endoderm staining. Mouse anti-CD56 (NCAM, 1:100) and rabbit anti-Brachyury (1:75) were used for Mesoderm staining. Alexa Fluor 488 anti-mouse (1:1000) and Alexa Fluor 594 anti-rabbit (1:1000) secondary antibodies were used as described above. Confocal microscopy analyses were performed as described above. All antibodies used are described in Table S2.

### Cytogenetic analysis

Conventional cytogenetic analysis was performed on 20 metaphase cells of 20 iPSC clones and parental AHF cells. Chromosomes were banded following standard methods for high-resolution G-banding. Cells were captured and karyotyped using a CytoVision Karyotyping System (Genetix, New Milton, UK).

### Differentiation into Neural Progenitor Cells

When cells reached 70-80% confluency, iPSCs were subjected to independent neuronal differentiations using STEMdiff Neuronal Differentiation Kit (STEMCELL Technologies). Per the monolayer culture protocol, single-cell suspensions were made for each clone to be plated, according to the manufacturer’s density suggestion, on a Matrigel-coated 6-well plate by dissociating cells with EZ Lift and replacing media with STEMdiff™ Neural Induction Medium containing Y-27632 (Rock inhibitor) and the corresponding supplement, SMADi. Daily media changes were performed for approximately 7 days with STEMdiff Neural Induction Medium and SMADi and returned to 37°C incubation overnight. Depending on the rate of cell growth, 80% of confluent wells were passaged in the range of 6-9 days post-plating. Mixed populations of neural progenitor and neural stem cells are present at this stage of differentiation and, therefore, were split for immunostaining and confocal analysis. Neural progenitor/stem cells were seeded onto 4-well or 8-well Matrigel-coated Nunc™ Lab-Tek II chamber slides according to densities recommended by the product manufacturer and adjusted based on the culture-ware surface area. Cells were maintained in STEMdiff Neural Induction Medium for 3-4 days to recover and attach to the surface at 37°C. Cells were washed with D-PBS without Calcium and Magnesium, fixed in PBS-2% PFA for 15 minutes, and permeabilized in PBS-2% PFA, 0.1% Triton X-100 at room temperature for 30 minutes. Cells were washed 3 times with D-PBS and incubated overnight in blocking solution (PBS-5% ES-FBS). Triple staining was performed using Anti-NESTIN Human, Anti-PAX6 and Anti-SOX1 antibodies and antibodies were incubated overnight a 4°C on a shaker. After three washes the corresponding Alexa Flour 488 mouse, Alexa Fluor 594 rabbit, Alexa Fluor 647 goat secondary antibodies were added for 2 hours incubation at room temperature. Slides were washed with PBS and cured overnight in Prolong solution containing DAPI and a glass coverslip. Confocal microscopy analyses were performed as described previously. All antibodies used are described in Table S2.

### Differentiation into Pancreatic Progenitor Cells

Pancreatic progenitor differentiation was performed using STEMdiff^TM^ Pancreatic Progenitor Kit (Stemcell Technologies) according to the manufacturer’s directions. iPSC cells were seeded in a single-cell suspension in a Matrigel coated 4-well chamber slide at a cell density of 4.5 x 10^5^ cells/cm^2^ in both a Matrigel-coated 12-well plate and 4-well chamber slide. Cells were differentiated over 12 days, during which a series of media changes containing six types of supplements were used corresponding to the day of differentiation to guide cells from an endodermal lineage towards a progenitor cell fate. By day 12, cells plated in the 4-well chamber slide were washed with D-PBS without Calcium and Magnesium (Gibco) and fixed in PBS-2% PFA for 15 minutes and permeabilized in 4% PFA 0.1% Triton X-100 at room temperature for 30 minutes. Cells were washed 3 times with D-PBS and incubated overnight in blocking solution (PBS-5% ES-FBS). Following overnight incubation, antibody mixtures were made with manufacturer-recommended dilution factors in a volume of blocking buffer (10% ES-FBS-PBS) prepared for each well. Triple staining was performed in distinct combinations for distinguishing markers associated with nuclear and surface proteins of progenitor (anti-SOX9, anti-PDX1, and anti-NKX6.1. Each antibody mix was added to the correct well and incubated overnight a 4°C on a shaker. Similar staining procedures were completed for the secondary antibodies with corresponding fluorophores for visualization: Alexa Flour 488, Alexa Fluor 594, Alexa Fluor 647 with an incubation period of 2 hours at RT. To finalize immunostaining, following five washes with PBS, the slides were cured overnight in Prolong solution containing DAPI (Cell Signaling Technology) with an adhered glass coverslip. Confocal microscopy analyses were performed as described previously. All antibodies used are described in Table S2

### Differentiation into Hematopoietic Progenitor Cells

iPSC clones were differentiated into Hematopoietic Progenitor Cells using the STEMdiff™ Hematopoietic Kit (Stemcell Technologies) according to the manufacturer’s directions. iPSC colonies were seeded on a matrigel coated 12-well plate according to the recommended density. Differentiation was performed for 12 days with different phases of media changes according to the manufacturer’s protocol. At day 10, 1 well of cells was harvested for a Colony Forming Unit Assay (CFU) and differentiated further utilizing MethoCult™ SF H4636 (Stemcell Technologies) for 14 days. Colonies from the CFU assay were counted and imaged between days 14 and 16 of the assay. The rest of the cells were harvested for flow cytometry at day 12. Cells were seeded at a density of 5×10^5^ cells per tube, blocked with a solution of PBS-5%ES-FBS-0.5% NaN_3_ at 4°C for 1 hour, immunostained using specific markers at 4°C for 1 hour, fixed with 2% PFA for 30 minutes, and washed with PBS. Triple staining was performed with Anti-Human CD43 Antibody PE (Stemcell Technologies), Anti-Human CD34 Antibody APC (Stemcell Technologies), and Brilliant Violet 421™ anti-human CD45 (BioLegend). Flow cytometry analysis was performed on a Cytek® Aurora flow cytometer, and analysis was performed using FlowJo (BD Pharminogen).

## Statistics

Initial data was processed in Microsoft Excel. The data were graphed using GraphPad Prism 9 software, and further statistical analysis was performed using the same software.

Reprogramming efficiency is presented as an average of at least 2, 3 or 4 independent experiments with at least 2 replicates. An unpaired t-test was used to analyze single comparisons between MeV and SeV reprogramming of each AHF and titration of MeV vector kinetics at each time point. Other quantitative data for all experiments are presented as the average of at least three independent experiments with at least two replicates for each experiment. All data are graphed as mean +/- SD. The statistical significance is specified as ∗p ≤ 0.05, ∗∗p ≤ 0.01. Comparisons not significant are not represented.

## Acknowledgment

We thank Dr. Kareem Mohni for reading the manuscript. We thank Qi Wang for technical assistance. This work was supported by Regenerative Medicine Minnesota (RMM 092319 DS 005 to P.D.), Mayo Graduate School, the National Institutes of Health (R56HL147852 and R01HL147852-A1 to P.D.), and National Resilience for financial support for this study. The graphical abstract was created using Biorender.com.

## Author Contributions

P.D. oversaw and conceptualized the project; P.D. designed and produced the vector construct; S.V., R.H., J.D., S.M., A.R., C.C., and B.S. acquired the data; P.D. and S.V. wrote the original first draft; P.D. reviewed, edited, and rewrote the manuscript that was approved by all authors.

---

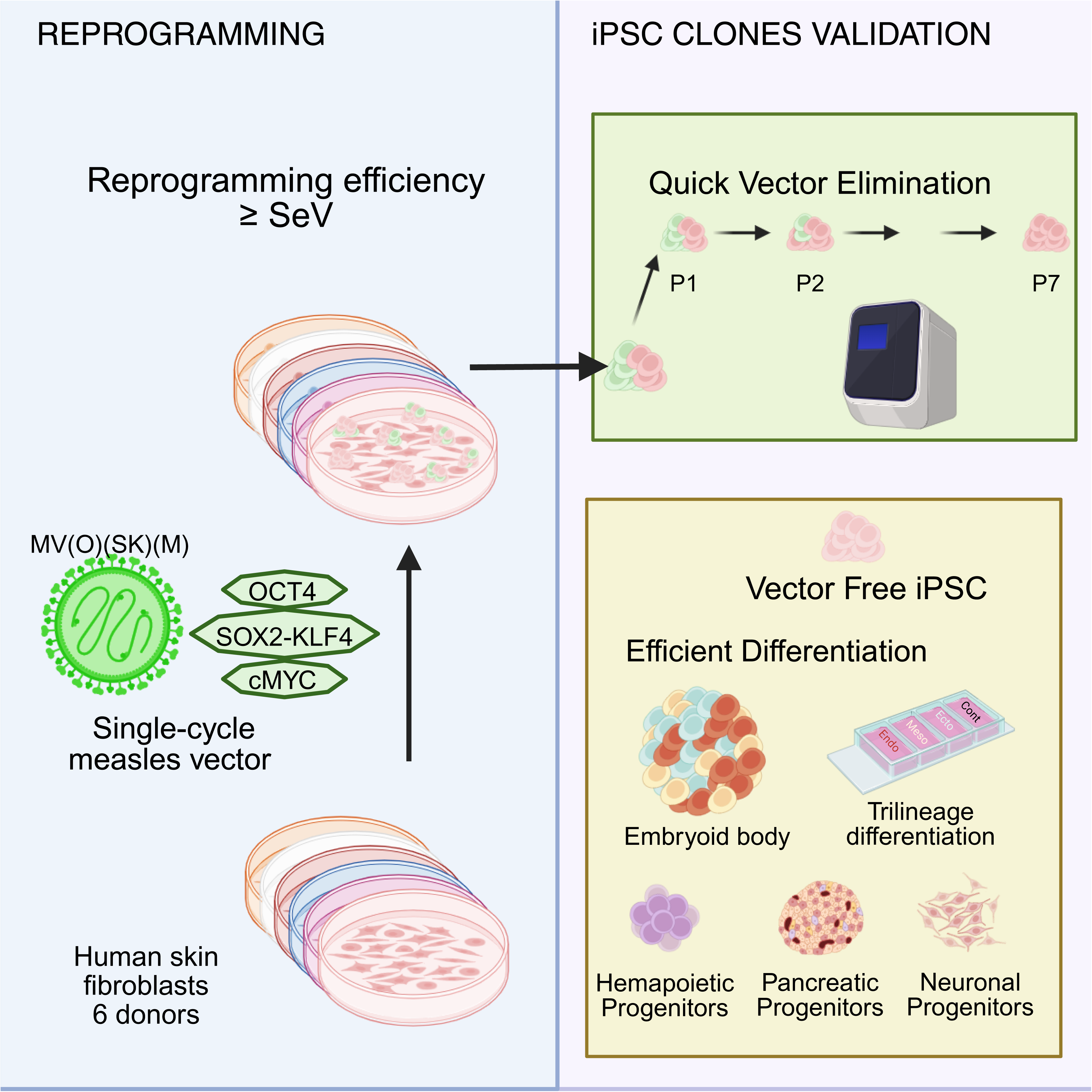

