## Supplemental data for "Efficient and reliable measles reprogramming platform for the generation of human iPSC"

**Table S1: Adult human fibroblasts**.

| Name | Male/Female | Age (y) | Passages reprogram |
|---|---|---|---|
| AHF1 | Male | 30 | p3, p4 |
| AHF2 | Male | 29 | p3, p4 |
| AHF3 | Male | 29 | p3, p4 |
| AHF4 | Female | 28 | p3, p4 |
| AHF5 | Female | 27 | p3, p4 |
| AHF6 | Female | 30 | p3, p4 |

**Table S2: Antibodies used for Immunofluorescence**

| PRIMARY ANTIBODIES | COMPANY | CAT |
|---|---|---|
| NANOG Rabbit Anti-human | Abcam | AB21624 |
| TRA-1-60 Mouse Anti-human (NovaRed) | Stemgent | 09-0010 |
| OCT4 Rabbit Anti-human | Cell Signaling Technology Inc | 2750S |
| SOX2 Rabbit Anti-human | Cell Signaling Technology Inc | 2748S |
| KLF4 Mouse Anti-human | Stemgent | 09-0021 |
| c-MYC Mouse Anti-human | Santa Cruz Biotechnology | sc-40 |
| mouse anti-SSEA-4 | Millipore | #SCR001 |
| mouse anti-TRA1-60 | Millipore | #SCR001 |
| mouse anti-TRA1-81 | Millipore | #SCR001 |
| rabbit anti-FOXA2 | Millipore | #07-633 |
| chicken anti-ß-III tubulin | Abcam | ab41489 |
| mouse anti-CD31 | Santa Cruz Biotechnology | sc-376764 |
| Rabbit anti-Nestin | Millipore | ABD69 |
| mouse anti-Pax-6 | Abcam | ab78545 |
| anti-SOX17 | R&D Systems | MAB1924 |
| rabbit anti-Brachyury | R&D Systems | MAB20851 |
| Anti-SOX1 Polyclonal Antibody | R&D Systems | AF3369 |
| anti-SOX9 | Abcam | ab185966-1001 |
| anti-PDX1 | Abcam | AB47383-1001 |
| anti-NKX6.1 | Bio-Techne | MAB5857 |
| Anti-Human CD43 Antibody PE | Stemcell Technologies | #60085PE |

| | | |
|---|---|---|
| Anti-Human CD34 Antibody APC | Stemcell Technologies | #60013AZ |
| Brilliant Violet 421™ anti-human CD45 | BioLegend | 304031 |
| **SECONDARY ANTIBODIES** | | |
| Alexa Fluor 647 Donkey Anti-Mouse | Life Technologies | A31571 |
| Alexa Fluor 594 Donkey Anti-Rabbit | Life Technologies | A21207 |
| Alexa Fluor 488 Donkey Anti-Mouse | Life Technologies | A21202 |
| Alexa Fluor 647 Goat Anti-Rat | Life Technologies | A21247 |
| FITC anti-chicken | Jackson ImmunoResearch | 703-095-155 |
| Alexa Flour 488 anti-goat | Jackson ImmunoResearch | 705-546-147 |

**Table S3: Primers for qPCR**

All sequences are human and written in the 5’ to 3’ direction with probe of 5’FAM and 3’TAMRA

| TARGET | FWD PRIMER | REV PRIMER | PROBE |
|---|---|---|---|
| *GAPDH* | ACCCAGAAGACTGTGGATG | TCAGCTCAGGGATGACCTT | CCCACAGCCTTGGCAGCGC |
| MeV-N | GGCCCAGCAGAGCAAGTGAT | TTGGCTGGACTCCGTTGCAG | AGCTGCCCATCTTCCAACCGGCA |

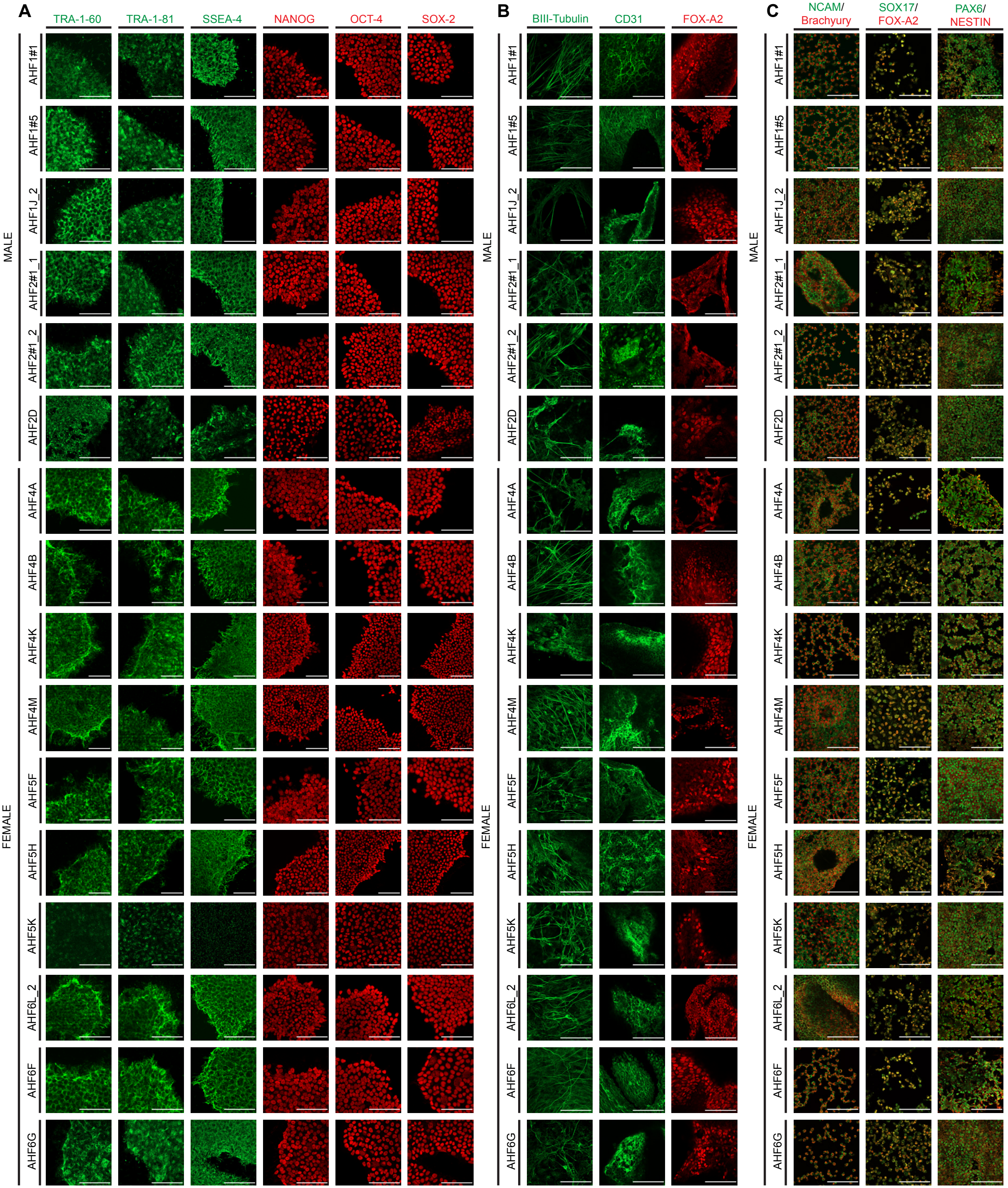
A
TRA-1-60
TRA-1-81
SSEA-4
NANOG
OCT-4
SOX-2
B
BIII-Tubulin
CD31
FOX-A2
C
NCAM/
Brachyury
SOX17/
FOX-A2
PAX6/
NESTIN
MALE
FEMALE
AHF1#1
AHF1#5
AHF1J_2
AHF2#1_1
AHF2#1_2
AHF2D
AHF4A
AHF4B
AHF4K
AHF4M
AHF5F
AHF5H
AHF5K
AHF6L_2
AHF6F
AHF6G

**Figure S1: MeV-derived iPSCs express pluripotency markers and can differentiate into the three germ layers.** (A) Confocal microscopy imaging from the 16 other MeV-derived iPSC clones with dual immunostaining for pluripotency markers TRA-1-60 (green)/NANOG (red), TRA-1-81 (green)/OCT4 (red), and SSEA-4 (green)/SOX2 (red). Scale bar: 100 µm. (B) Confocal microscopy imaging of spontaneous differentiation of 16 other MeV-derived iPSC clones into embryoid bodies (EB). Cells were immunostained for EB markers βIII-TUBULIN (green), CD31 (green), and FOX-A2 (red). Scale bar: 100 µm. (C) Guided trilineage differentiation into mesoderm, endoderm, and ectoderm germ tissues of 16 other MeV-derived iPSC clones. Cells were immunostained for associated lineage markers using NCAM (green)/BRACHYURY (red) for Mesoderm, SOX 17 (green)/FOXA2 (red) for Endoderm and PAX6 (green)/NESTIN (red) for Ectoderm lineages. Scale bar: 100 µm

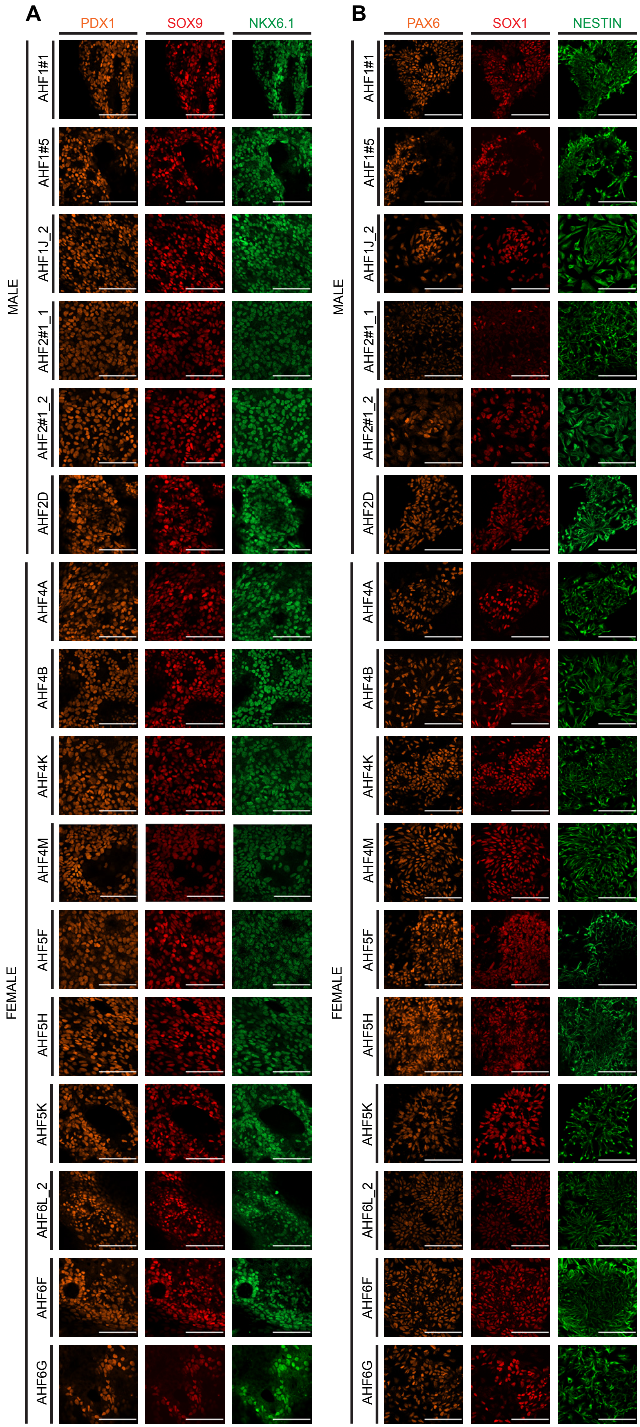

A
PDX1
SOX9
NKX6.1
B
PAX6
SOX1
NESTIN
MALE
FEMALE
AHF1#1
AHF1#5
AHF1J_2
AHF2#1_1
AHF2#1_2
AHF2D
AHF4A
AHF4B
AHF4K
AHF4M
AHF5F
AHF5H
AHF5K
AHF6L_2
AHF6F
AHF6G

**Figure S2:** (A) Confocal microscopy imaging of neuronal progenitor differentiation from the 16 other MeV-derived iPSC clones. Cells were stained with PAX6 (orange), SOX1 (red), NESTIN (green), and DAPI (blue). Scale bar: 100 µm (B) Confocal microscopy imaging of pancreatic progenitor differentiation from the 16 other MeV-derived iPSC clones. Cells were immunostained for PDX1 (orange), SOX9 (red), NKX6.1 (green), and DAPI (blue). Scale bar: 100 µm.
